# Non-Optimal Codon Usage Regulates Cell Cycle Progression: Functional Insights Of Codon Optimization of CDK1 and NUF2 Genes

**DOI:** 10.1101/2025.11.05.681855

**Authors:** Mahua Bhattacharya, Eliezer Gideon Baum, Milana Frenkel Morgenstern

## Abstract

The expression and functions of genes are largely dependent on genome integrity and stability. Codon usage plays a significant role in maintaining stability and functions of genes. Perturbation in codon sequence can lead to functional and structural dysfunction of a gene. In our study, we performed the codon usage analyses of cell cycle-dependent genes. Various codon usage parameters analyzed using nucleotide compositions like RSCU, ENC, and GC analyses showed that cell cycle dependent genes follow non-optimal codon usage. Genes preferred AT-rich ending over GC-rich ending, thereby suggesting preference for non-optimal codons. Neutrality and parity plot showed that the codon preference is a result of mutation selection pressure. To study the codon usage implications, we optimized the codons of cell cycle dependent genes (CDK1 and NUF2) to study their effects on cell cycle and apoptosis *in vitro*. We observed that codon optimization alters the cell cycle length in cell cycle–dependent genes impacting cell fate and survival. Our studies revealed that codon usage preferences directly affect the stability of both mRNA and proteins. Specifically, genes and proteins with non-optimal codons exhibited reduced stability compared to their optimized counterparts, suggesting critical implications for cell cycle regulation and apoptosis.

## INTRODUCTION

There are 20 amino acids in the mammalian system that constitute the formation of proteins. These amino acids are coded by three nucleotides called codons. In humans, there are 64 codons, of which 3 codes for stop codons and other 61 codes for amino acids. Except for methionine and tryptophan, all other amino acids are coded by more than one codon ^1,2^. The preference of selecting one codon over another is called codon usage bias (CUB). Synonymous mutations are mutations where changes in nucleotide does not change the amino acid^3^. Most amino acids are coded with more than two codons and the frequency of using the codons is attributed to CUB. Synonymous mutations may alter the mRNA stability, co-translation protein folding and time of elongation during translations ^4–7^.

CUB is studied using various nucleotide compositions, where the CU is a frequency of an effective number of synonymous codons, and it is different for every species. CUB (CUB) is reflecting a mutational or selection pressure. CU and CUB are largely depending on GC content, gene expression, gene length, nucleotide composition, tRNA abundance, hydropathicity and aromaticity of proteins^8–10^. In humans, the “optimal” codons are the ones which have either a G or C in the third position^10^. Other parameters that are used to study codon bias are RSCU, ENC, Codon Adaptation Index (CAI), and GC3 content. RSCU is the ratio of observed frequency of synonymous codon to frequency of all codons for those amino acids multiplied by degeneracy levels. RSCU values of >1.6 are overrepresented while <0.6 are underrepresented for a given codon. While frequently used codons are >1, less frequently used codons are <1 and codons that are randomly selected are often represented as 1 ^11,12^. ENC is the effective number of codons for a given amino acid. In a gene or length of genome ENC is inversely proportional to CUB, which means if a gene has higher ENC, it prefers one codon that is much higher than the other. ENC and GC3 are usually calculated to observe the biasness of codons for different amino acids^8,9,13^. CAI is also a measure of CUB. CAI reflects the extent to which specific codons are preferentially adapted relative to synonymous alternatives. ^13^.

The study of genetic code has been done widely to study the effect of CUB on various cellular and phenotypic mechanisms. CUB is also widely used to study virus-host interaction. Many of the RNA viruses have non-optimal codons to the host^14^. This helps viruses to integrate and infect. Carmi G *et.al*, studied CU of SARS-Cov-2 and identified that it uses non-optimal codons, which helps to infect a plethora of hosts^14^. CU of HPV suggests that the non-optimality of HPV to human genome helps in integration of virus to human genome^15^.

CUB has been associated with genes related to obesity. Study done by Chakraborty S *et.al*, suggested that obesity genes have low CUB compared to the housekeeping genes^8^. Similarly, they have also studied the association of CUB on genes associated with central nervous systems. The study showed that genes that are associated with diseases like dementia, Alzheimer’s, have significant usage of low CUB. The neutrality and parity analysis showed that mutation selection pressure might be involved for low CUB^9^.

CU patterns have also been studied in human embryonic stem cells. Modified tRNA at inosine bind with wobble effect. The abundance of the tRNA causes increased translation speed of self-renewing genes while stalling the translation speed of genes involved in differentiation. This regulates the pluripotent stem cells self-renewal and differentiation mechanism ^16^. CU have also been studied in maintaining the mRNA stability and controlling the transcription mechanism during meiosis^17^.

A study by Frenkel-Morgenstern et al. demonstrated that cell cycle–dependent (periodic) genes preferentially use non-optimal codons. Their analysis highlighted the role of aminoacyl-tRNA synthetases (aaRS), showing that WARS (preferred codon) and GAPDH (constitutive control) maintained stable expression throughout the cell cycle, whereas TARS and GLARS displayed cell cycle–dependent expression patterns (Figure 4).^18^. The study also suggests that the expression of aaRS across the cell cycle is also dependent on availability of tRNA and affinity of a codon for tRNA.

In this study, we have studied the codon usage bias between cell cycle dependent genes and non-cell cycle dependent genes. We optimized two genes viz., *CDK1* and *NUF2* to study the effect of optimization on stability and functions of these genes. We identified that optimization of these genes affects their stability both on mRNA and protein levels. We observed changes in cell cycle and apoptosis as a result of codon optimization. Therefore, it is observed from our study that cell cycle dependent genes use non-optimal codons.

## Materials And Method

### i. Codon Usage Analysis

All Refseq-ids were converted to gene-ids using the R-package biomaRt. The coding sequence of each gene was extracted from the GRCh38-p13 CDS file obtained from the ensemble website using *in-house* Python script. The longest isoform for each gene was selected to avoid bias. The human codon usage table was obtained from the kazusa database (http://www.kazusa.or.jp/codon/). Then the COUSIN software (V1.0) was used to calculate all the nucleotide statistics which include: A3, T3, G3, C3, GC-all, GC1, GC2, GC3, ENC and several other codon-usage indices. Then t-tests were conducted between both the Periodic and Non-periodic groups (two tail t-tests). Finally, using R, a parity-plot, GC12 - GC3 plot and ENC-GC3 plot (GC12 was calculated as a mean between GC1 and GC2 for each gene) were calculated.

### ii. RNA Sequencing and DEG

Transcriptomic datasets were obtained from public datasets (https://www.ncbi.nlm.nih.gov/gds). GSE145372, GSE122512 for cervical cancer transcriptome data was downloaded for DEGs for comparative analysis with 4C seq data. GSE94479 and GSE104736 for HeLa DEG and MCF-7 for DEG in different cell cycle phases for codon usage study were downloaded. Differential gene expression analysis was performed using EdgeR. The Benjamini–Hochberg method for controlling the false discovery rate (FDR) was used for adjusting p-values. Genes with FDR < 0.05 and fold-change ≥ 2 were considered as differentially expressed.

### iii. Gene ontology analysis

Gene ontology analyses were performed using PantherDB^19^ and Metascape^20^. Pathway analysis was done using KEGG and the Reactome pathway database.

### iv. Molecular Cloning

The genes of *CDK1, CDK1op, NUF,* and *NUFop,* were synthesized by Twist Biosciences with AgeI and NheI restriction sites. The plasmid pEGFP-N1 and genes were double restricted and digested using AgeI-HF (New England Biolabs.inc cat no: R3552) and NheI-HF (New England Biolabs.inc cat no: R3131). 1µg of plasmid was used for digestion and the reaction was carried out using manufacturer’s protocol.

The mixture was incubated for 1 hr at 37℃, followed by heat inactivation at 65℃ for 15 mins. The digested product (plasmid and genes) was run on 1% Agarose gel and band of appropriate sizes were cut and purified using Nucleospin Gel and PCR purification kit (cat no: 740609.50).

The Digested genes were ligated to digested plasmid using T4 DNA ligase (New England Biolabs incat no: M0202S). The ligation was carried out using the manufacturer’s protocol.

A molar ratio of 1:3 was considered for ligation. The ligation mixture was incubated O/N at 16℃. Heat inactivation was carried out at 65℃ for 10 mins.

### v. Transformation

The ligated products were transformed in DH5α competent cells. 100µl of competent cells were thawed on ice. The 2 µl of ligated product was added to the competent cells’ tubes. Heat shock at 42℃ was given for 90 sec. immediately after heat shock the tubes were transferred on ice for 5 mins. 1ml LB broth was added to the cells and incubated at 37℃ for 45 mins. The competent cells with plasmid were streaked on LB agar plates with 100µg/ml of Ampicillin. The plates were incubated O/N at 37℃. Next day, the colonies were picked and incubated in 1ml LB broth each with 100µg/ml of Ampicillin. Plasmids were isolated using Nucleospin Plasmid Mini Kit® (cat no: 740588.50). The plasmids were sent for sanger sequencing. Colonies with confirmed sequencing results were used to make glycerol stocks for the plasmids and used for plasmid isolations later.

### vi. Transfection

pEGFP-N1 CDK, pEGP-N1 CDKop, pEGFP-N1 NUF, pEGFP-N1 NUFop, was transfected in HEK293T, SiHa, HeLa and C33a using Polyjet™ In-Vitro Transfection reagent (SignaGen laboratories, cat no: SL100688) as per manufacturer’s protocol. Plasmid to polyjet was used in 1:3 ratio for HEK293T while for HeLa, SiHa and C33a were used in 1:4 ratio, incubated for 15 mins and added to the cells in cell plate. After 6hrs post transfection, the media was replaced with fresh media and cells were observed for GFP after 24hrs under fluorescent microscope.

### vii. *in-silico* RNA Folding

*in-silico* RNA folding pattern was studied using Vienna RNAfold Package 2.0^21^. The sequence of the genes was uploaded directly on the webserver. The parameters were kept default. Minimum fold energy was calculated for each gene. The MEFs determine the stability of the RNA structure.

### viii. *in-vitro* RNA Stability Assay

*In-vitro* RNA stability assay was performed using actinomycin D. Hek293T cells were cultured in 6-Well plates. The cells were transfected with pEGFP-N1 CDK, pEGP-N1 CDKop, pEGFP-N1 NUF, pEGFP-N1 NUFop, pEGFP-N1. 24hrs post transfection, cells were treated with 10µg/ml of Actinomycin D. The cells Were collected at 0, 4, 9, 16 and 24hrs as per *Liu.Y et.al* ^22^. RNA of the cells was extracted using Qiagen® RNeasy mini purification kit (cat no: 74004). The purified RNA was eluted in RNAse and DNAse free water.

### ix. qPCR

Extracted Actinomycin D treated RNA was quantified using nanodrop. Approximately 500ng of RNA was converted to cDNA using High-Capacity cDNA Reverse Transcription Kit (cat no: 4368814) using manufacturer’s protocol. For qPCR, SYBR® Green qPCR from Thermo Fischer Scientific SYBR Green Real-Time PCR Master Mixes ™ was used to perform qPCR as per manufacturer’s protocol. Each mixture was made in triplicates for each gene and GAPDH was used as internal control. NTC was used as control.

The primer for qPCR was designed using Primer3Plus (https://www.primer3plus.com/index.html)

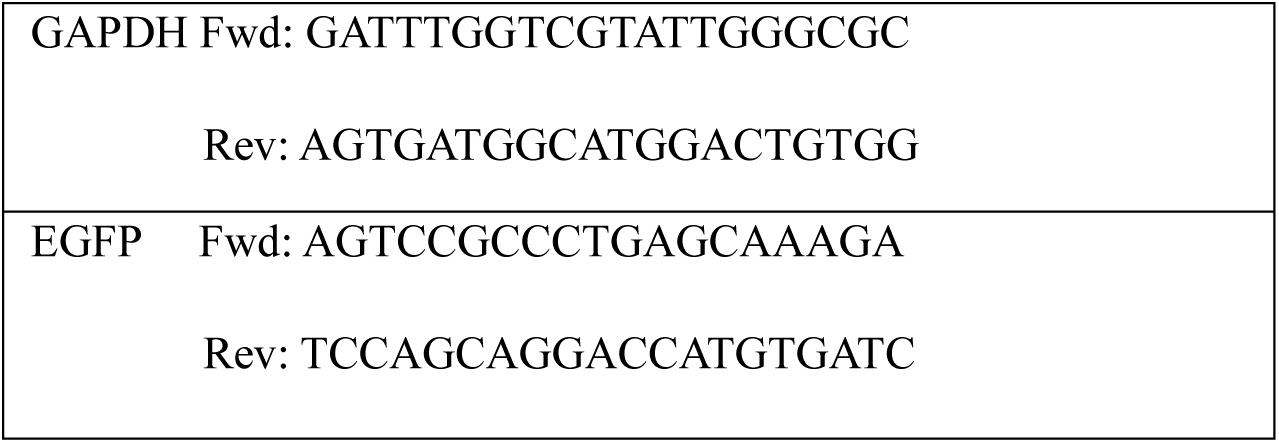

Standard qPCR was performed. Analysis was done using NTC as control and GAPDH as relative expression control. 2^−ΔΔct^ was calculated against 0 hr actinomycin D treated RNA.

### x. *in-vitro* Protein Stability Assay

Thermal shift assay was performed to study *in-vitro* protein stability^23^. HEK293T cells were transfected with pEGFP-N1 CDK, pEGP-N1 CDKop, pEGFP-N1 NUF, and pEGFP-N1 NUFop. 24hrs post transfection, cells were harvested, and protein was isolated using RIPA buffer (see appendix for recipe). 20µg of protein lysate was subjected to various temperatures 4℃, 42.3℃, 44.6℃, 47℃ and 52℃ for 3 mins and transferred on ice for 5 mins. A 5X loading buffer (see appendix for recipe) was added to the samples for Western blot analysis. The samples were run on SDS-PAGE and transferred on Nitrocellulose membrane (Tamar). Anti-GFP antibody was used in 1:8000 as primary antibody (Abcam cat no: AB-ab183734) and Goat anti-Rabbit IgG (Abcam AB-ab6721) as secondary in 1:20000 dilutions and visualized under chemiluminescence ChemDoc. The bands were quantified using ImageJ software.

### xi. Cell Cycle Analysis

HEK293T, HeLa, C33a and SiHa were transfected with pEGFP-N1 CDK, pEGP-N1 CDKop, pEGFP-N1 NUF, pEGFP-N1 NUFop, for 24 hrs. Post transfection, transfected cells were trypsinized and washed with PBS 2X times. Cells were fixed with 1% formaldehyde for 15 mins at 37℃. After fixation, cells were washed with PBS 3 times and permeabilized with 0.25% triton X100 (Sigma) for 15 mins at RT. Cells were washed with PBS and suspended in 1ml PBS. Hoechst 33342 (Sigma-Aldrich) was used at 3µg/ml concentration and cells were incubated in dark at RT for 15mins. Cells were analyzed in MoFlo Astrios Flow cytometer (Beckman-Coulter). Untransfected and unstained cells were used as negative control. The gating was done for GFP. The cells with GFP were further analyzed for Hoechst33342 stain for cell cycle analysis^24^. Analysis was done using Flow-Jo analyzer.

For synchronization, cells were transfected. Post 24hrs of transfection, cells were treated with double thymidine block^10,24,25^ which arrests the cells in G0/G1 phase. Cells were subjected to thymidine at 100µg/ml concentrations. Cells were released from synchronization at 0^th^, 6^th^, 10^th^, 13^th^ and 15^th^ hr. Cell cycle analysis was performed using Hoechst 33342 like the above protocol for non-synchronized cells.

### xii. Apoptosis

HeLa, C33a, SiHa were transfected with pEGFP-N1 CDK, pEGP-N1 CDKop, pEGFP-N1 NUF, pEGFP-N1 NUFop. 24hrs post transfection cells were trypsinized and stained with AnnexinV-7AAD as per manufacturer’s protocol. Cells were suspended in 1X binding buffer. 5 μL of the Annexin V-CF Blue and 5 μL of 7-AAD were added to each 100 μL of cell suspension. Cells were incubated at RT for 15 min in the dark. After incubation, 400 μL of 1X Annexin-binding buffer was added to the cell suspension and proceeded for flow cytometry. Untransfected cells, heat-treated cells and unstained cells were used as control. Flow cytometry analysis was performed for apoptosis on Galios (Beckman-Coulter).

## RESULTS

### 1. Identification Of Cell Cycle Dependent Genes from Transcriptome Data

Cell cycle phase transcriptomic datasets of HeLa, U2OS and MCF-7 cell lines were downloaded from public dataset (GEO accession no: GSE94479 and GSE104736)^26–28^. The differential gene expression analysis was performed using EdgeR software (see Methods) on these dataset and common gene clusters were used for this study (Figure 1A) with *p-value* <0.01 and an FDR of <0.05, the gene set of differentially expressed genes were divided into two sets viz., periodic (expression of the genes dependent on cell cycle and non-periodic (the gene expression is not dependent on a cell cycle). 259 genes were classified as periodic genes, while the rest were classified as non-periodic. Gene ontology analysis was performed on periodic dataset using Metascape ^20^. The gene ontology analysis suggests large cluster of genes involved in cell cycle, regulation of cell cycle process, cell cycle checkpoints and cell cycle etc., highlighted in green. (Figure 1B).

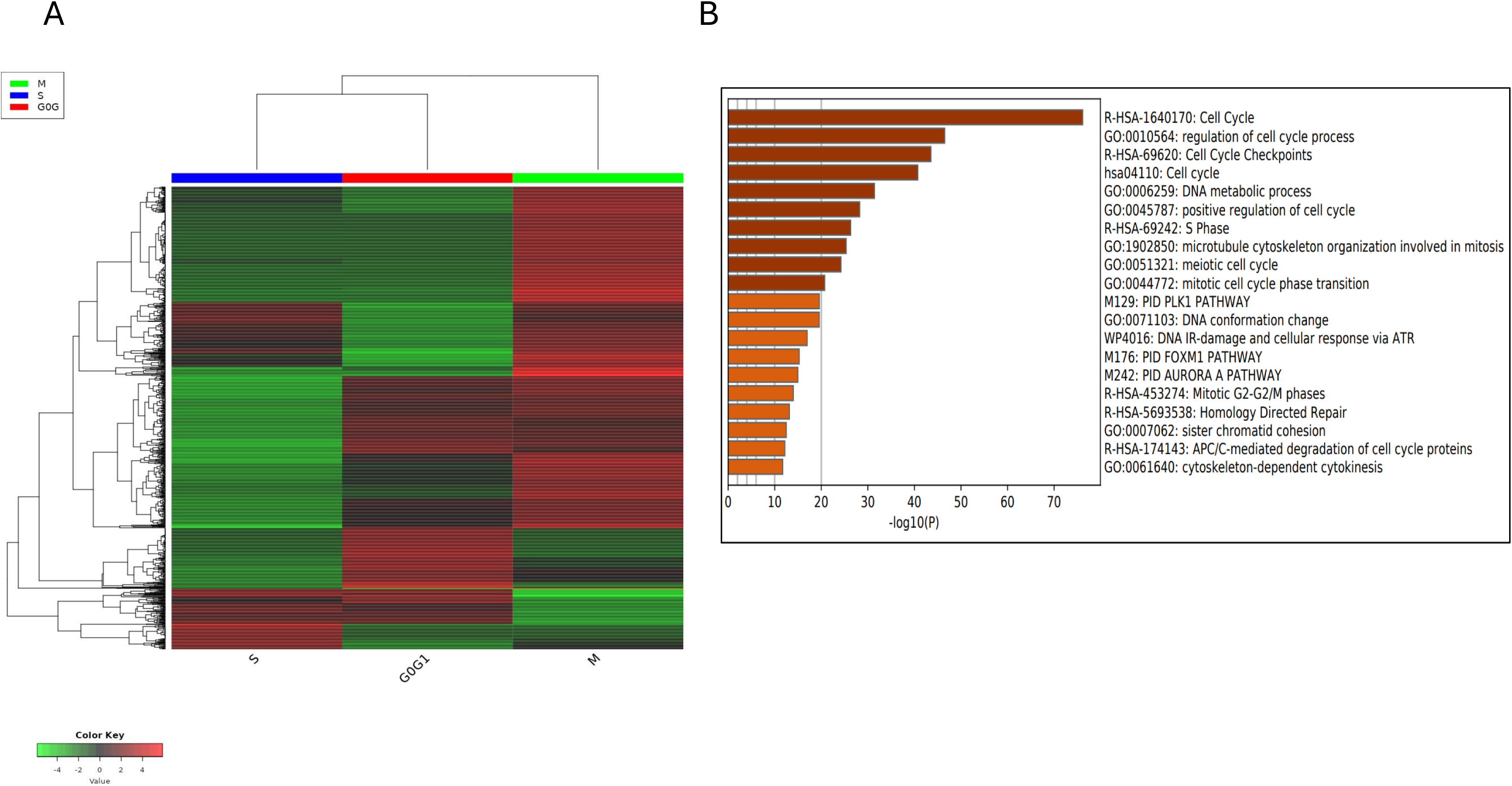

Next, to study the CU patterns, as Well as RSCU, CUSP, CAI, ENC measurements, neutrality and parity analyses were performed on both periodic and non-periodic genes. These parameters help to understand the codon bias patterns of the genes. To study the biases, only the longest isoform of transcripts was taken into consideration for both datasets.

### 2. The Nucleotide Composition Analysis

Different nucleotide position analysis was done between periodic and non-periodic genes (Table 1). The results suggest that overall GC3 content of periodic genes is less compared to non-periodic genes. Periodic genes have more A and T at the 3^rd^ position than the non-periodic genes. The G3 and C3 positions also suggest that periodic genes have less G and C at 3^rd^ position compared to non-periodic genes. Other CUB analyses such as COUSIN, CAI, SCUO are used to calculate overall bias. The analyses show that the codon usage of periodic genes is non-optimal compared to non-periodic genes. The GCall study which refers to overall GC content shows periodic genes use less GC than non-periodic at *p-value <0.05.* These analyses suggest that periodic genes are non-optimized compared to periodic genes. The statistical analysis using two tailed t-test with p-value >0.05 proved that the difference between nucleotide composition between periodic and non-periodic genes are statistically significant. These results indicate that periodic genes use non-preferred codons (non-optimal codons) over non-periodic genes which use more optimized codons.

**TABLE 1:** Statistics for the various nucleotide position and nucleotide composition between cell-cycle-regulated and non-cell-cycle-regulated groups of genes.

| Stat | T | Pvalue | Periodic | Non |
| --- | --- | --- | --- | --- |
| A3 | 5.375 | 1.86E-07 | 22.223 | 18.740 |
| T3 | 4.082 | 6.14E-05 | 23.772 | 21.381 |
| G3 | -3.162 | 1.77E-03 | 27.623 | 29.276 |
| C3 | -5.722 | 3.22E-08 | 26.382 | 30.602 |
| GCall | -4.767 | 3.29E-06 | 49.943 | 52.853 |
| GC1 | -2.337 | 2.03E-02 | 54.865 | 55.967 |
| GC2 | -3.646 | 3.28E-04 | 40.960 | 42.712 |
| GC3 | -4.993 | 1.17E-06 | 54.005 | 59.879 |
| COUSIN_18 | -4.839 | 2.38E-06 | 0.585 | 1.251 |
| COUSIN_59 | -4.884 | 1.94E-06 | 0.567 | 1.089 |
| CAI_59 | -4.959 | 1.37E-06 | 0.750 | 0.767 |
| CAI_18 | -5.544 | 8.02E-08 | 0.783 | 0.801 |
| ENC | 2.256 | 2.50E-02 | 48.453 | 47.435 |
| SCUO | -2.168 | 3.12E-02 | 0.126 | 0.135 |
| FOP | -4.741 | 3.70E-06 | 0.512 | 0.569 |
| CBI | -4.908 | 1.73E-06 | 0.064 | 0.176 |
| ICDI | -1.990 | 4.77E-02 | 0.199 | 0.216 |
| Scaled_Chi | 3.222 | 1.45E-03 | 0.422 | 0.397 |
| GRAVY | -6.559 | 3.40E-10 | -0.501 | -0.344 |
| AROMA | -5.505 | 9.65E-08 | 0.070 | 0.079 |

### 3. The RSCU Analysis and Codon Frequency Analysis

First, the frequency of codon was analyzed for periodic genes. The codon frequency analysis (Nc) suggests the frequency of observed codons in a set of genes. Nc suggests that cell-cycle-regulated (or periodic) genes use non-optimized codons (Figure 2A). However, I also observed that frequency of CTG was high compared to other codons. Next, RSCU of periodic genes was analyzed. The RSCU value was calculated for all periodic genes. The RSCU value of periodic genes was to be <1.0 which means that the codons were less frequently used. Most of the periodic genes were concentrated at the center of the axes. Axis1 and Axis2 occupied nearly 38% and 26% respectively. RSCU analysis for cell cycle dependent genes showed that most of the codons are non-optimized and optimized codons are used less frequently (Figure 2B).

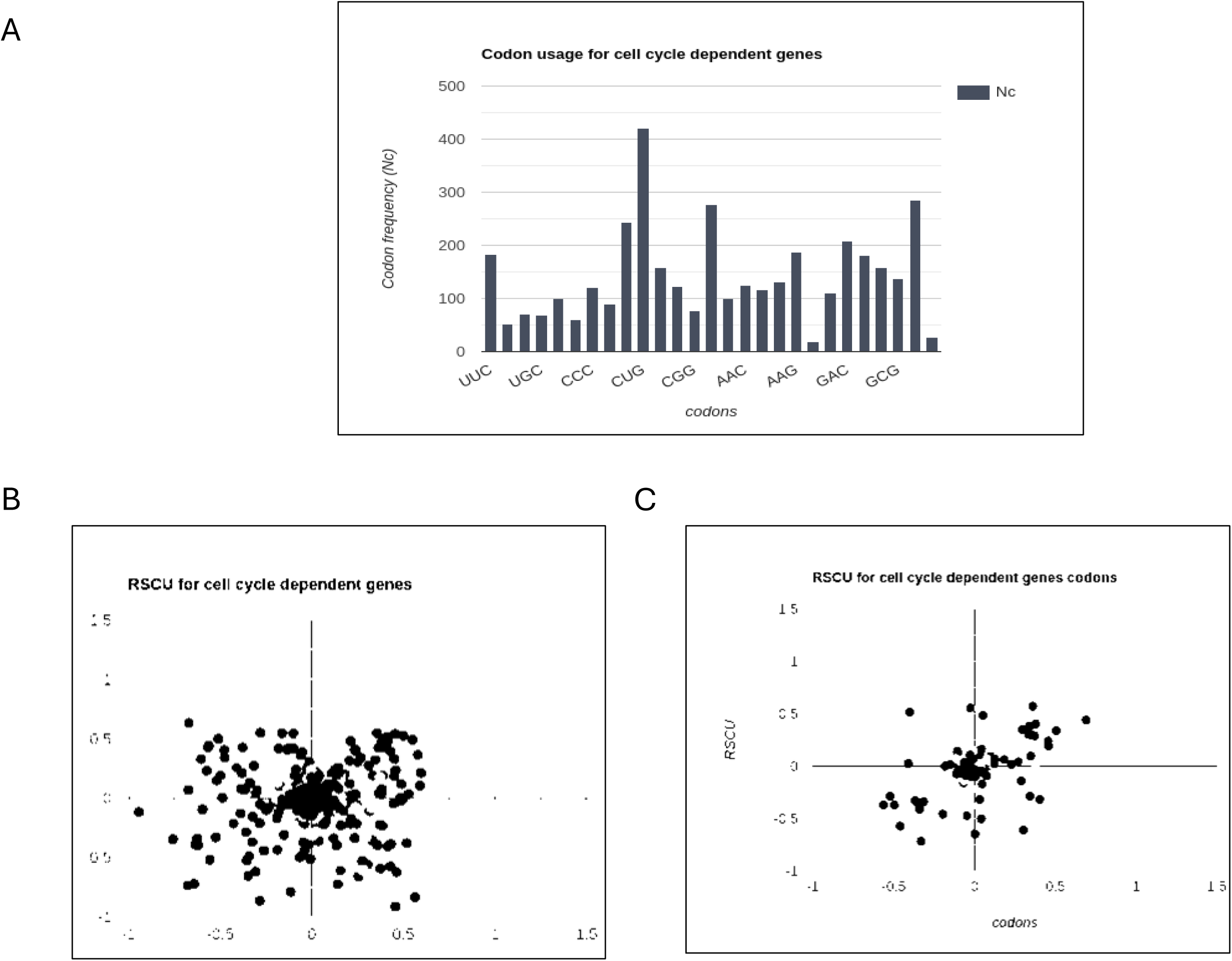

### 4. ENC, neutrality and Parity analysis

ENC is an effective way of calculating the CUB. The correlation between ENC and GC3 was investigated. I observed that ENC-GC3 correlation was higher in non-periodic genes compared to periodic genes (Figure 3A). This result indicates that most of the non-periodic genes prefer optimized codons while periodic genes are biased to non-optimized codons.

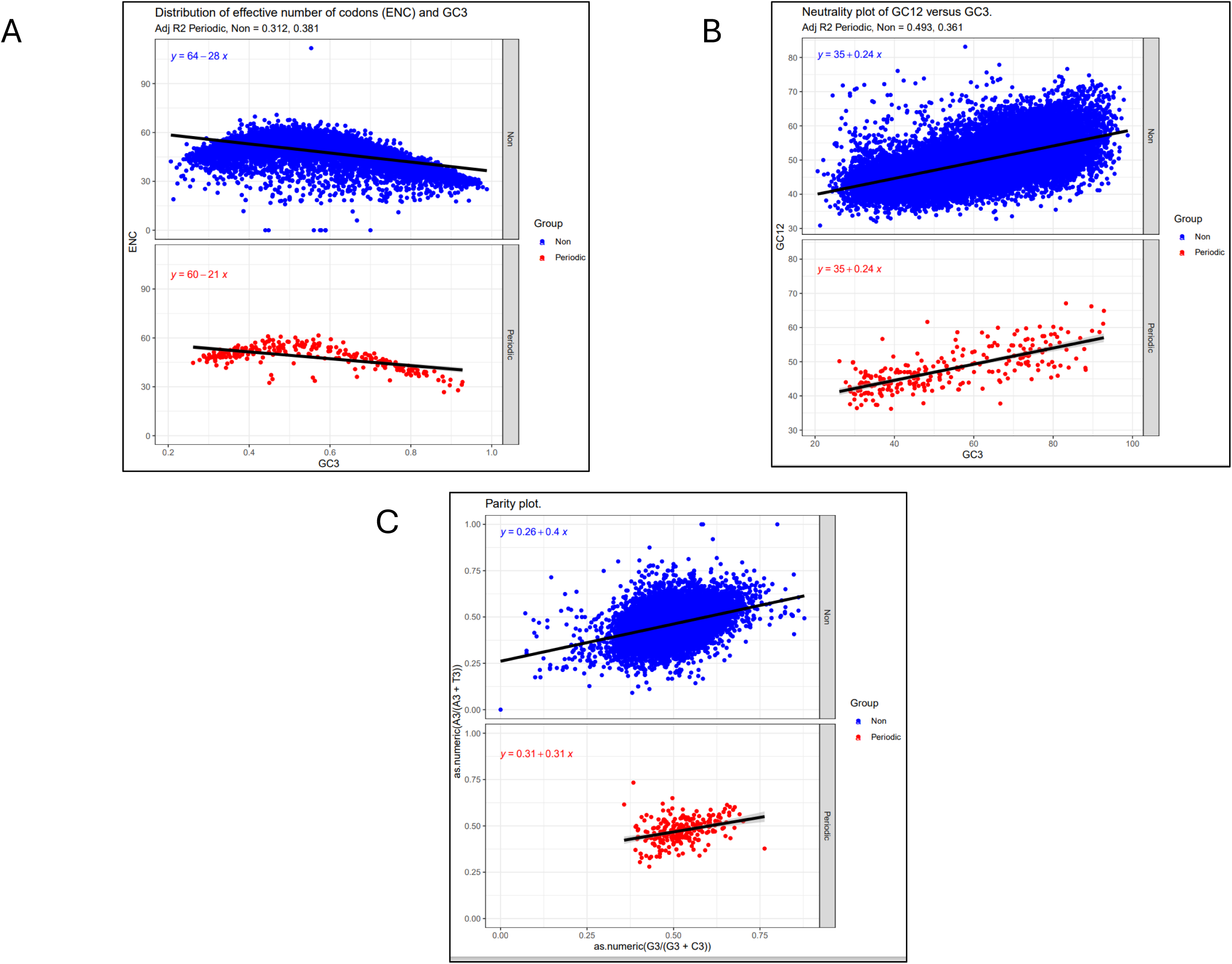

Next, the neutrality and parity analysis were performed. This analysis helps in understanding natural vs mutational selection. Neutrality plot is used to study the association of GC3 to GC12. If the correlation is significant, it implies that CUB was a result of mutational selection. However, if the correlation is not significant, it implies that selection is purely under natural selection pressure^8^. In my study, I found the correlation between GC12 and GC3 of periodic and non-periodic genes was significant with *p-value*<0.05 (Figure 3B). This indicates that the CUB was under mutational selection pressure.

Parity plot helps in understanding the correlations between purine and pyrimidine. The distribution of codons across A3/T3 and G3C3 was calculated for periodic and non-periodic. I observed that distribution of codons across A3-T3 and G3-C3 was not even. The non-periodic group showed distribution across the linear axis while periodic genes were more clustered 0.4 to 0.75, thereby not showing distribution across the linear scale (Figure 3C). This suggests that CUB could be a consequence of mutational selection rather than natural selection.

### 5. Optimization Of Cell Cycle Regulated Genes

To study the effect of codon optimization on cell cycle dependent genes, I selected three genes viz., *CDK1, NUF2 and ARLIP6I*. For optimization, CAI was taken into consideration. CAI close to 0.99-1 means optimized codons. The codons of these three genes were optimized using JCat^29^. The CAI value of genes before and after codon optimization can be seen in (Figure 4) and (Table 2) The genes were then cloned *in-house* modified pEGFP-N1 vector. The reporter EGFP was chosen as it has mammalian optimized codons which will prevent any form of bias that would be caused by non-optimized codon GFP for downstream analysis. The genes were cloned in the vector at NheI and AgeI site followed by C-terminal EGFP. The cloned plasmids were then transfected in HEK293T, HeLa, SiHa, CaSki and C33a cell lines using polyjet for various experiments.

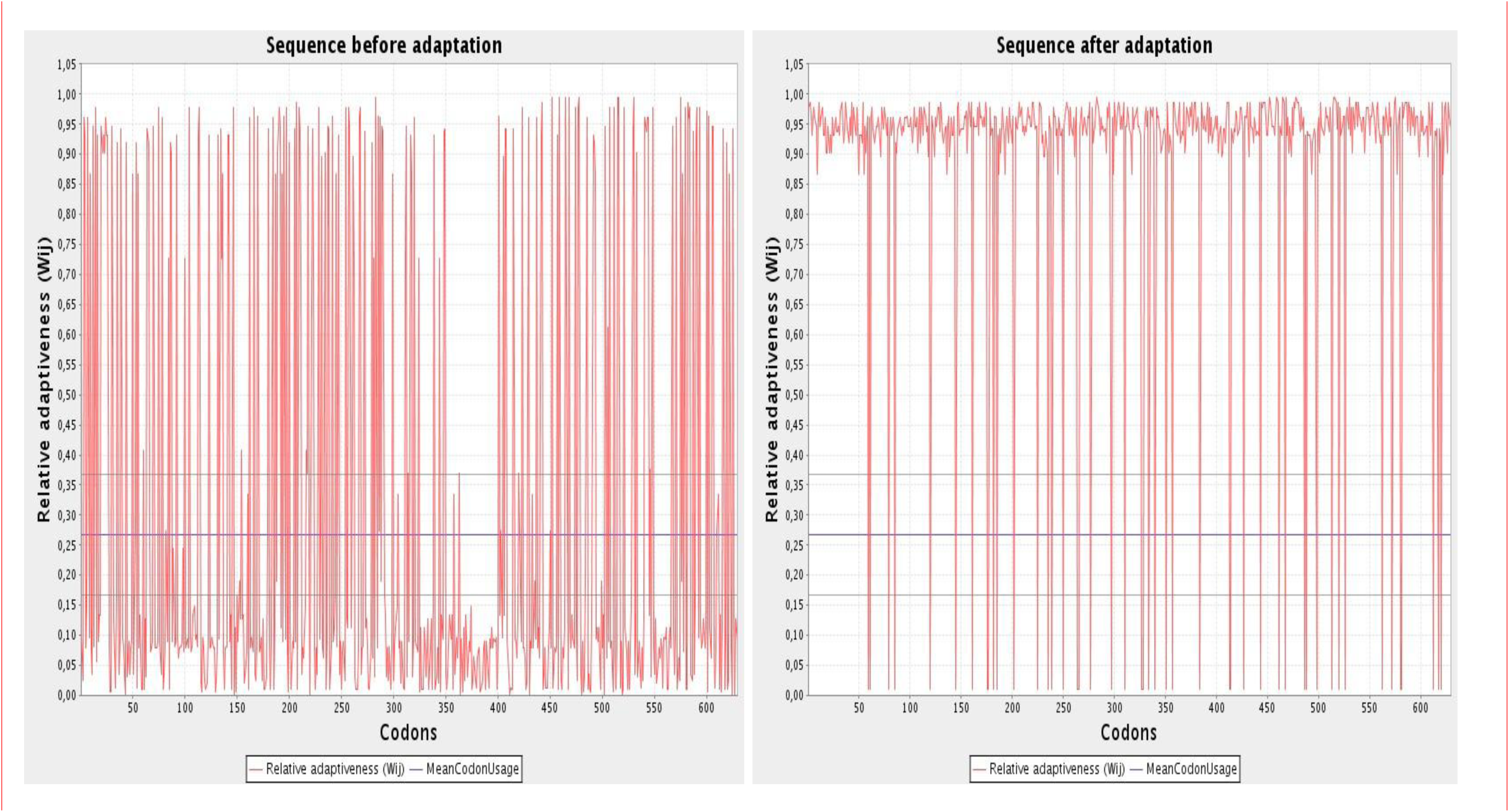

**TABLE 2:** CAI values of three genes before and after optimization.

| Genes | CAI | Optimised CAI |
| --- | --- | --- |
| CDK1 | 0.63 | 1.000 |
| NUF2 | 0.675 | 1.000 |

Codon optimization affects mRNA stability, protein stability, and protein misfolding, thereby affecting the overall structure and functions of the genes. In our study, we employed *in-silico* and *in-vitro* analysis of mRNA stability. Protein misfolding and the effect of codon usage bias on translational rate can be studied using thermal shift assay, trypsin-sensitive assay, and cycloheximide assay.

### 6. Effect of Codon Usage Optimization on RNA Stability

To check the effect of optimizing codons on RNA, RNA stability using *in-silico* webserver, namely, Vienna RNAfold^21^ was performed. mRNA folding plays an important role in mRNA stability^30,31^. The minimum fold energy was calculated to determine the fold pattern and stability of the mRNA. Lower MFE is associated with stable RNA structure while higher MFE is associated with relatively unstable structure. From *in-silic*o analyses of fold pattern between non-optimized and optimized sequence of all six genes, MFE for all six genes were calculated. We observed that MFE of optimized genes is lower than that of the non-optimized (Figure 5A-B).

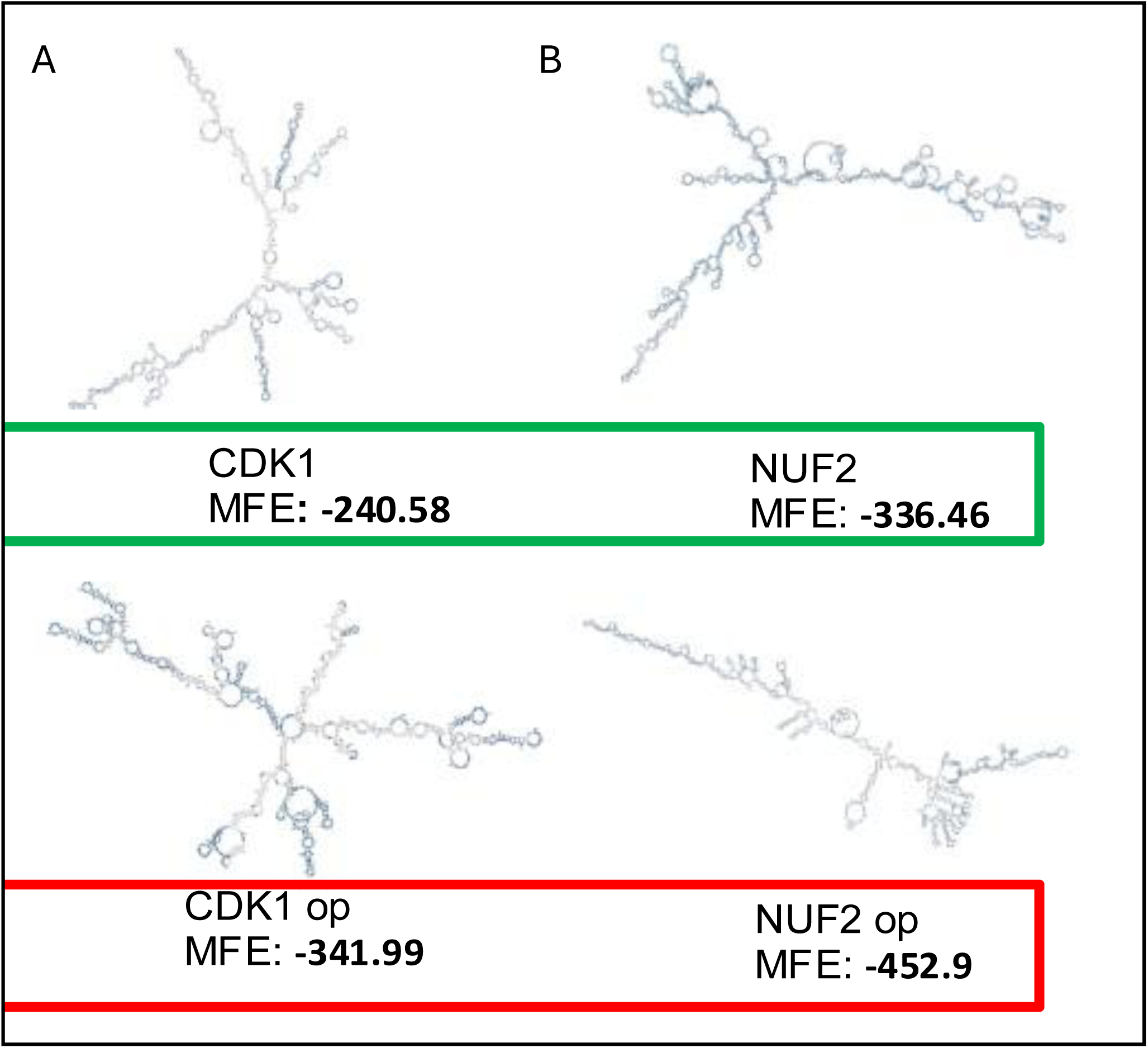

To validate the *in-silico* prediction of mRNA stability, *in-vitro* mRNA stability was performed using Actinomycin D (10µg/ml)^22^. HEK293T cells were transfected with the plasmids of optimized and non-optimized genes for 48 hrs. The cells were treated with Actinomycin D (10µg/ml), a transcriptional inhibitor post-transfection. The treated cells were collected at different time-points and RNA was isolated. Real-time PCR was performed on these mRNA and against *GAPDH* as internal control. The mRNA levels of *CDK* decreased from the 9^th^ hrs significantly (*p-value <0.05*), while mRNA levels of *CDKop* were significantly stable. Similar results were observed for *NUF2* and *NUF2op*. The difference in mRNA levels between optimized and non-optimized *CDK* and *NUF2* were statistically significant with *p-value <0.05.* The results suggest that optimized genes were stable in the system and that this result proves the *in-silico* results (Figure 6).

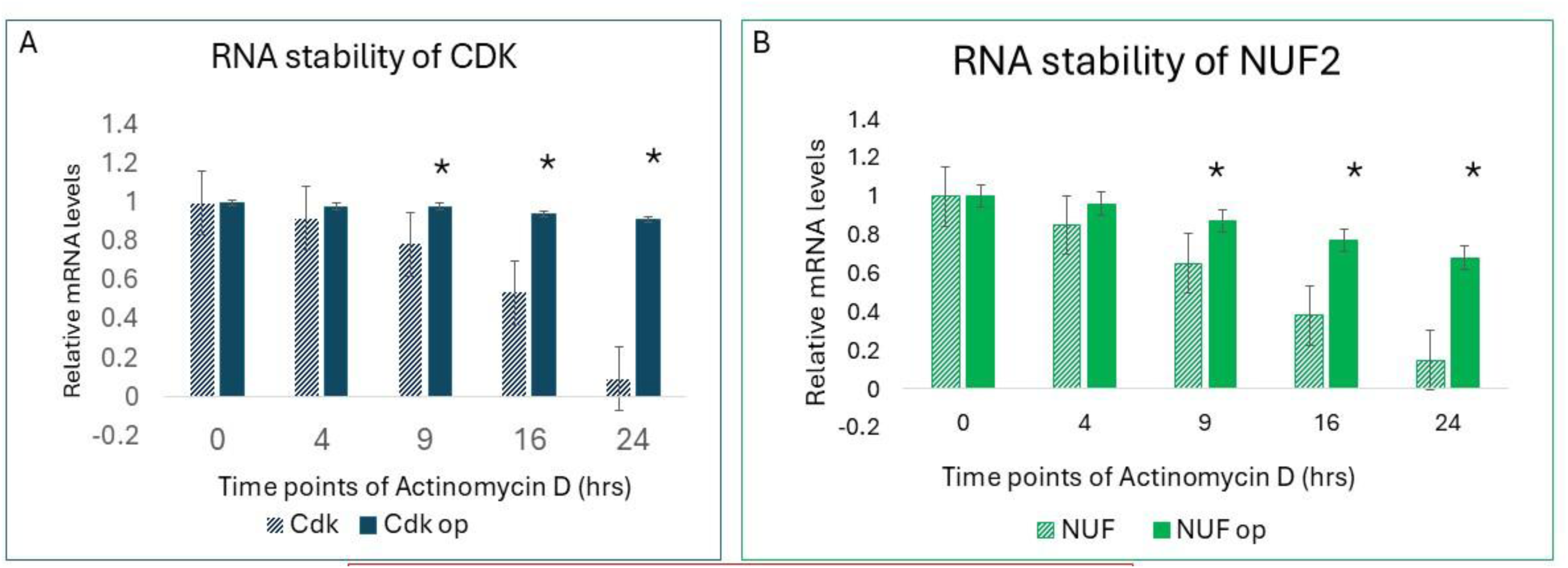

### 7. Effect Of Codon Optimization on Protein Stability

Next, to check if codon bias has any implication on protein stability, protein stability assay was performed *in-vitro*^32^. HEK293T cells were transfected by pEGFP-N1-*CDK/ CDKop/ NUF/ NUFop* for 24hrs. For thermal shift assay the transfected cells were harvested and lysate were subjected to different temperatures for 3mins and immediately cooled on ice for 5 mins. Western blot was performed with anti-GFP antibody. I observed that protein stability was different for all the genes. CDK1 was stable until 44.6℃, it started losing its stability beyond 47℃ (*p-value <0.05*). However, CDK1op was relatively stable and maintained its structure throughout the temperatures. A similar pattern was observed for NUF and NUFop at *p-value <0.05* (Figure 7A-B). These results indicate that non-optimized CDK1 and NUF2 are more stable than CDK1op and NUF2op.

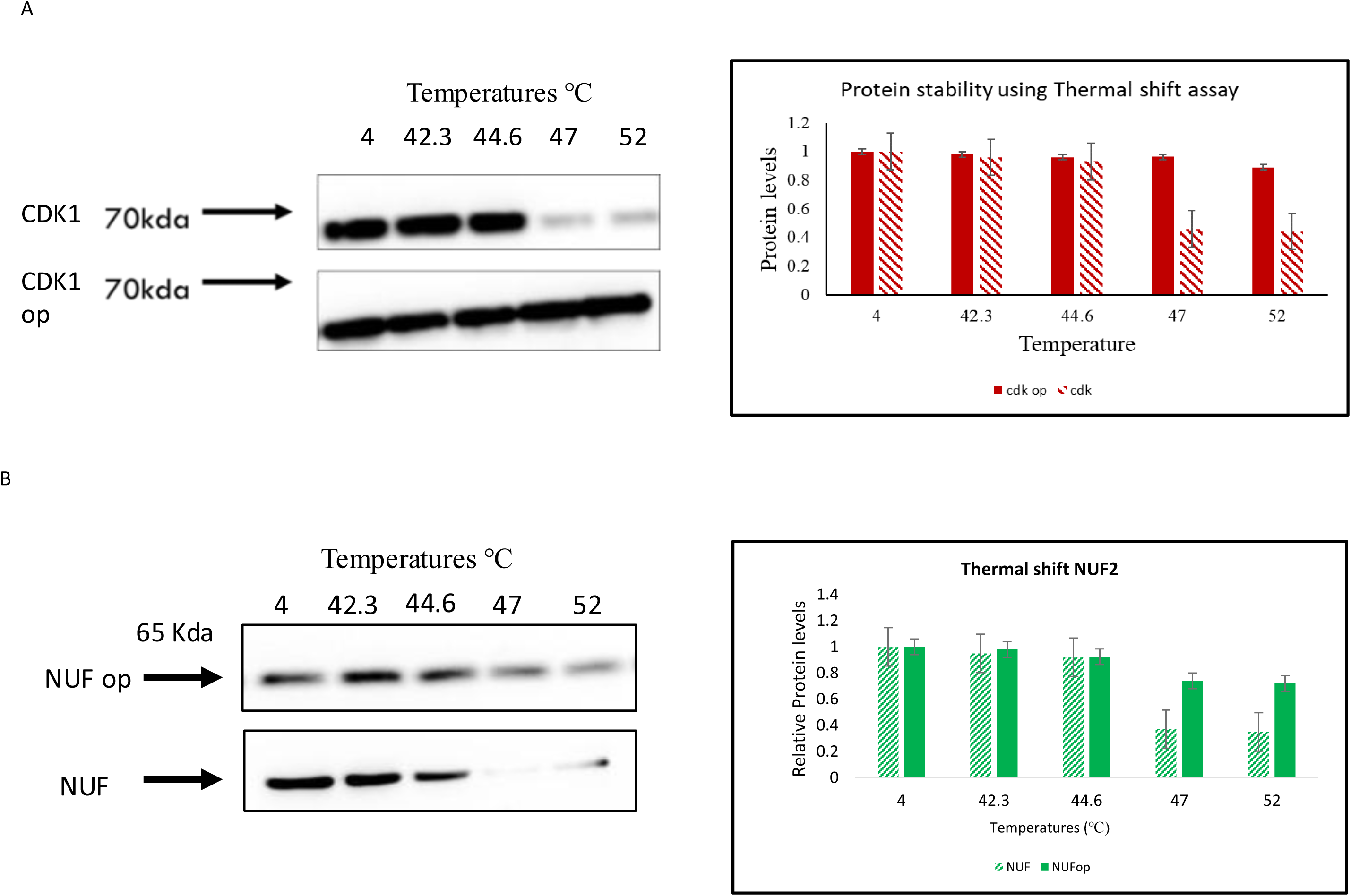

### 8. Effect of codon optimization on cell cycle and apoptosis

#### A. Cell cycle

After stability, next we studied the effect of synonymous CU on functions of cell cycle dependent genes. We transfected and overexpressed both optimized and wildtype genes on HEK293T cell lines and performed cell cycle analysis on asynchronized cells. Gating was done using untransfected cells and gated as GFP-ve (Figure S1A). Next, the transfected cells were gated for GFP+ve. Approximately 50000 were used for analysis. GFP +ve cells were gated for cell cycle analysis. The gated GFP +ve cells were further gated for cell cycle phase. The gating for G0/G1, S and G2/M phase was done using Hoechst stain using UV laser in flow cytometer (Figure S1B). The experiment demonstrated that optimized genes enter G2 phase earlier than non-optimized genes (Figure 8).

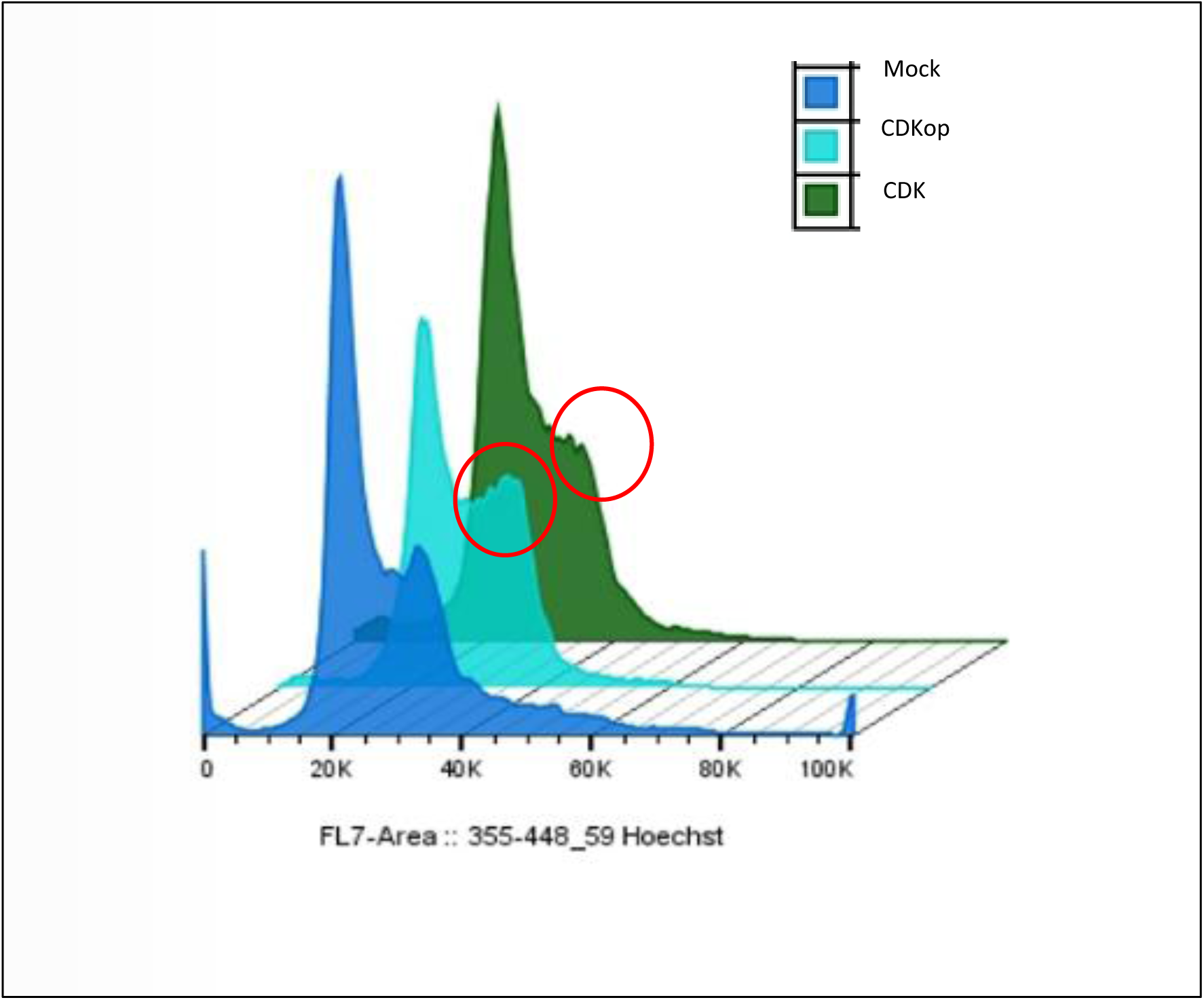

To elucidate further, synchronization was performed on HEK293T cell line transfected with optimized and non-optimized CDK1 genes. After 24 hrs, cells were treated with double thymidine blocks and released. The released cells were fixed and stained with Hoechst. We observed that genes with optimized codons enter G2 phase earlier than non-optimized genes (Figure 9). To elucidate the results further, we performed synchronization of transfected cells using double thymidine block. CDK1, CDK1op, NUF2 and NUF2op plasmid construct was transfected in HeLa, SiHa and C33a. The transfected synchronized cells were released at different time points and analyzed the cell cycle phase using flow cytometer. The statistics were performed using *in-built* statistics of FlowJo®. Flow cytometric analysis suggested that cells with CDK1 and NUF entered G2 phase later compared to CDK1op and NUFop cells. CDK1 and NUF transfected cells were observed to enter G2 at 15^th^ hrs while CDK1op and NUFop cells entered cell cycle at 10^th^ hrs (Figure 10A-B). These results indicate that non-optimized genes of CDK, and NUF have longer cell cycle length compared to optimized CDKop, and NUFop which have shorter cell cycle length.

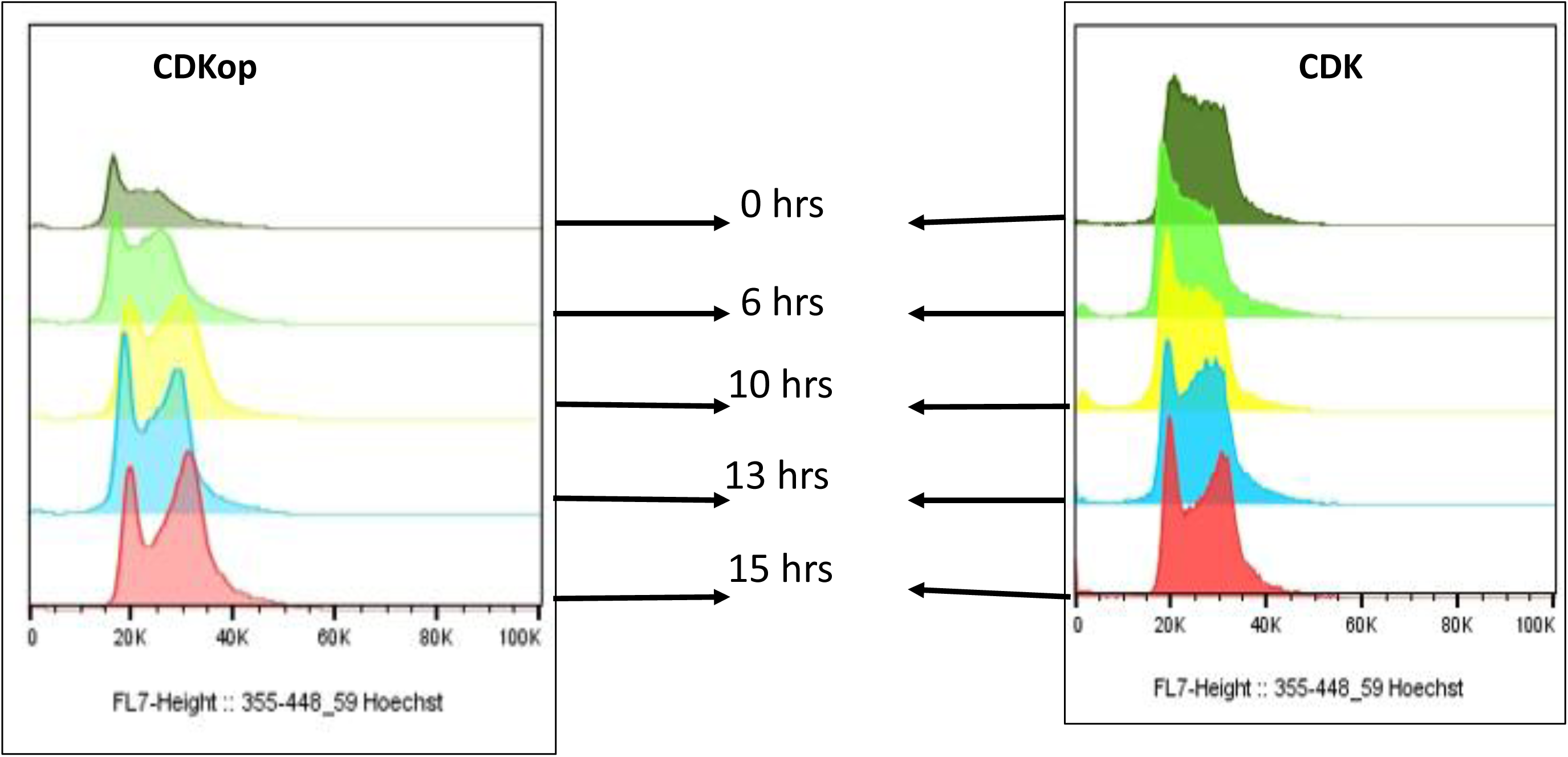

**Figure 10.**
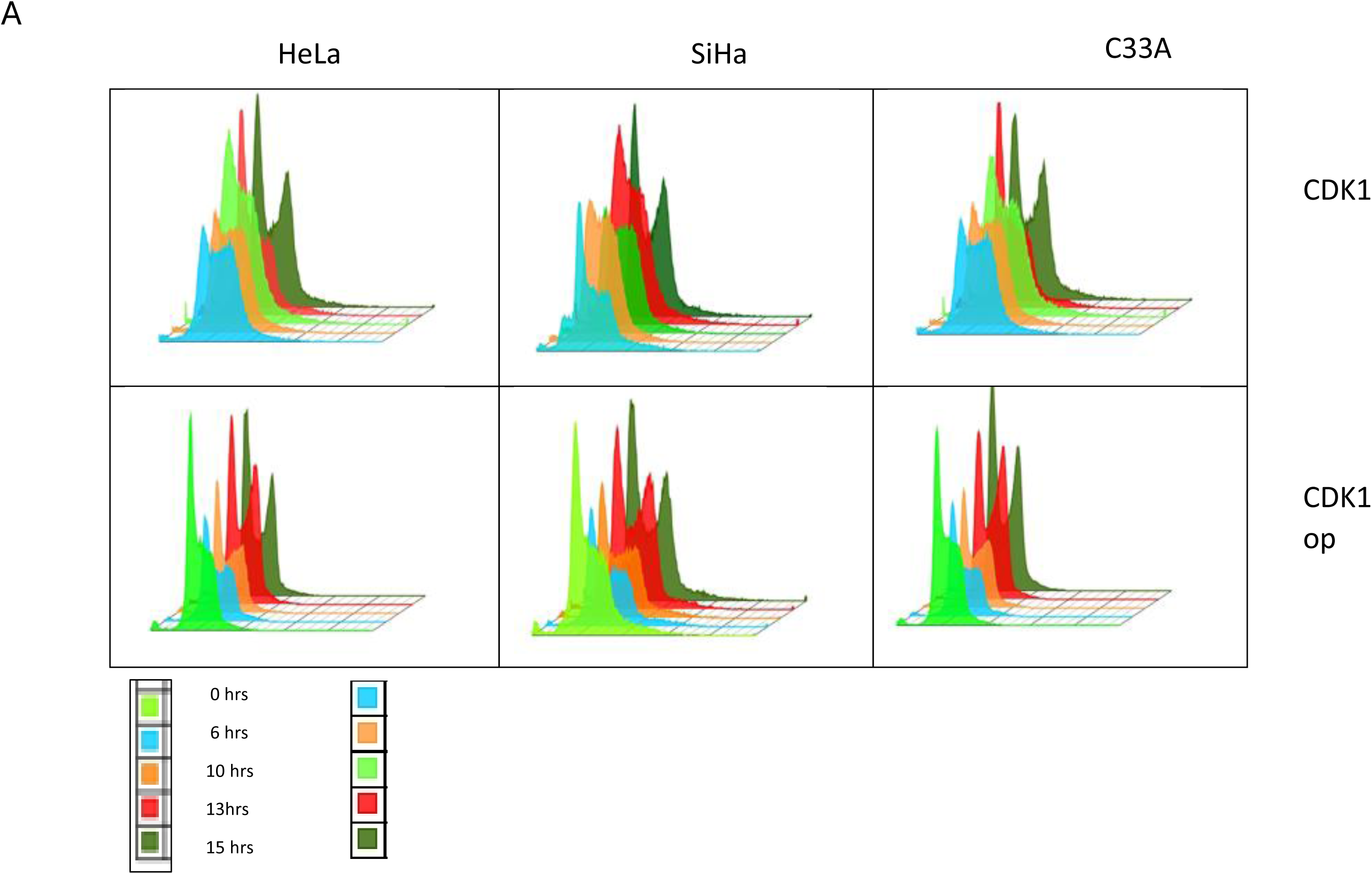
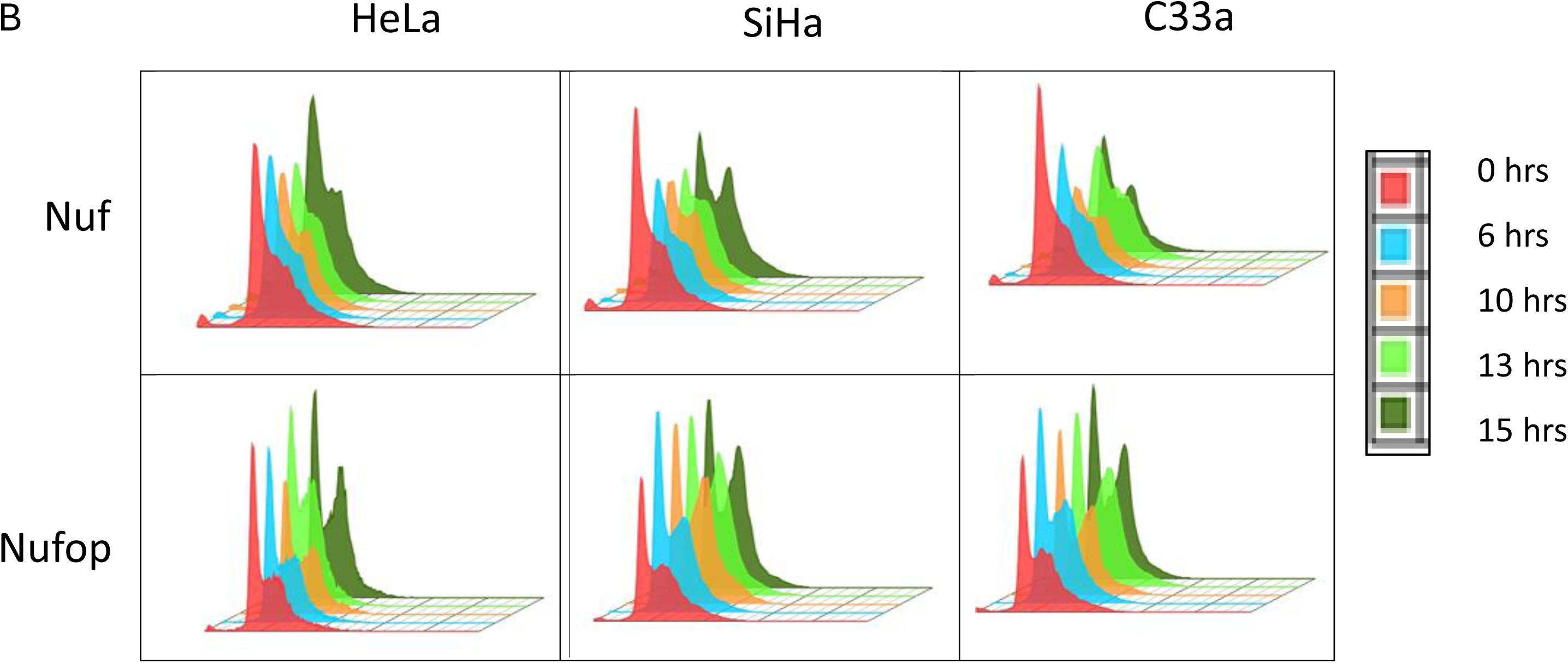
Flow cytometry analysis of Cell cycle synchronized cells. A: Top: CDK1 (L-R): HeLa, SiHa and C33a cells enter G2 at 15^th^ hr **Bottom:** CDK1op (L-R): **B: Top:** NUF (L-R): HeLa, SiHa and C33a cells enter G2 at 15^th^ hr **Bottom:** NUFop (L-R): HeLa, SiHa and C33a cells are observed to enter G2 at 10^th^ hr. (*p-value <0.05)*

#### B. Apoptosis

Next, we checked for the effect of codon optimization on apoptosis. SiHa, HeLa and C33a cell lines were transfected with respected plasmids. After 24 hrs the cells were harvested and stained with Annexin V and 7AAD. Approximately, 50000 cells were used for analysis (Supplementary Figure S2).

Apoptosis analysis of CDK and CDKop showed that CDKop cells had undergone more apoptosis than CDK in all three cell lines at *p-value<0.05.* Similarly, NUF and NUFop showed similar results (Figure 11). Apoptosis analysis of cells transfected with optimized and non-optimized codon proved that optimized genes transfected cells undergo apoptosis at a much higher rate compared to non-optimized transfected cells.

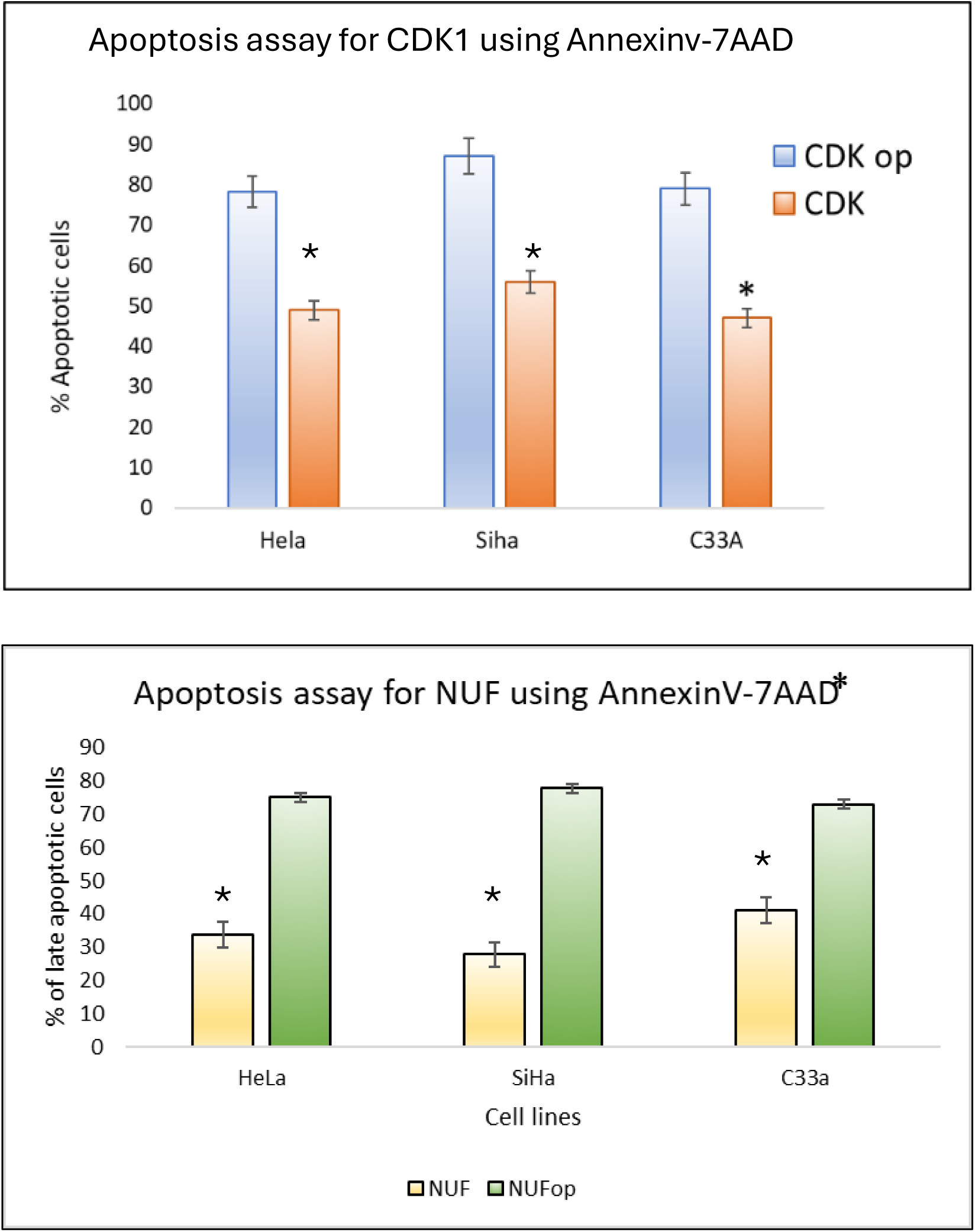

The results indicate that codon optimized genes shorten the cell cycle, but the cells later undergo apoptosis. This phenomenon could possibly be associated with “mitotic catastrophe”. Our studies indicate that optimization of codons plays a role in cell survival and cell fate. However, further analysis needs to be done to understand the exact mechanism of how cells with optimized codons undergo apoptosis.

## DISCUSSION

Synonymous mutations have not been explored completely in cancer or other genetic disorders^33,34^. Synonymous mutations have been associated with genes in RNA splicing^35,36^. Studies of synonymous mutations have been done on HNSCC^37^, pancreatic^38^ and ovarian cancer^39^.However, studies on synonymous mutations have been linked to understanding the CU preference of a gene. CUB plays an important role in defining mRNA stability^40^, protein folding and protein stability^41,42^.

In my study, CUB was elucidated in cell cycle dependent genes. RSCU analysis performed on cell cycle dependent genes suggested that most of the cell cycle dependent genes involved in cell cycle process prefer non-optimal CU with RSCU score <1. Human genes have a tendency of preferring G/C codons over A/T ending codons^36^. The analysis of ENC and GC3 of these genes suggested that cell cycle dependent genes have preference for low frequency codons, also called non-optimal codons.

My study on understanding the selection process of the codons, whether natural or mutational selection pressure caused codon bias, neutrality and parity analysis was performed. This analysis states that if the GC12 vs GC3 correlates, then the selection is a result of mutational selection pressure. Similarly for parity if the correlation is significant then the codon bias is a result of mutation selection^8,9^. In my study I found the correlation for both neutrality and parity statistically significant, thereby suggesting that the CUB of cell cycle dependent genes is probably skewed towards non-optimal codon usage, because of mutational selection pressure.

CU has been linked to mRNA stability, translational efficiency and altered mRNA splicing structure^43–46^. The study on *E.coli* and *S.cerevisiae* showed that the rate of translation was codon dependent. These organisms tend to prefer AT ending codons over GC. When the codons were changed to non-optimized i.e., GC ending, the rate of translation was slower than the optimized codons^47,48^.

Codon usage plays an important role in RNA stability^17,22,30,43,49^. RNA stability and RNA folding play an important role in studying gene expression analysis as well as translation efficiency^30,50^. The stability of mRNA secondary structure holds a positive correlation with mRNA abundance. ^30,50–52^ Greater stability of mRNA results in greater protein abundance due to efficient translation and high fidelity^30,31,49,50,52,53^. An *in-silico* analysis of RNA stability was performed. The MFE of non-optimized was observed to be higher than optimized. This suggested that optimized genes were more stable than non-optimized. Further validation was done using *in-vitro* mRNA stability assay^22^. HEK293T cells were transfected with non-optimized and optimized codon genes. The cells were treated with Actinomycin D and qPCR was performed with *GAPDH* as internal control. I found that genes with non-optimized codon (*CDK1* and *NUF2*) were less stable compared to genes with optimized codons (*CDK1op* and *NUF2op*). The degradation of mRNA was observed from the 9^th^ hr for non-optimized as compared to optimized that was relatively stable.

The translation efficiency is also attributed to availability of tRNA and ribosome stacking^14^. In non-optimized codons, ribosome stacks and thereby slows the rate of translation. This rate and speed of elongation determine protein folding and protein stability^42,54^. After mRNA stability, protein stability assay was performed to understand the stability of proteins. A thermal shift assay was performed on transfected HEK293T cells^32^. The cells were harvested, and protein was subjected to different temperatures for 3 mins. My study showed that non-optimized proteins were less stable and degraded beyond 44.6℃ while optimized proteins were stable. However, from previous studies, it was shown that KRASop was less stable than dopKRAS. CU of KRAS impacts its function on transcription. Optimized KRAS had altered transcriptional activity because of its effect on histone acetylation and methylation^22^. This study led to the understanding that the translational speed is responsible for the misfolding of proteins. Since, optimized genes have high speed of transition and elongation rate due to no phenomena of ribosome stacking, leading to unstable and misfolded proteins^22,55^. However, a study on FRQ protein in *neurospora* showed that optimized FRQ was more stable than dopFRQ and an increased opFRQ levels were observed than dopFRQ. However, opFRQ resulted in perturbed function of FRQ. It was suggested that FRQ, which has intrinsically disordered regions, helps in the binding of other proteins for functioning of FRQ. Therefore, despite opFRQ being stable had functional implication^23^. CU can affect the drug target binding site seen in *MDRI* gene^56^.

After performing the stability assay for mRNA and proteins for the genes, functional implication of codon optimization was studied. Since our study was on cell cycle dependent genes, we wanted to elucidate the role of non-optimized and optimized genes of *CDK1,* and *NUF2* on cell cycle. Three cervical cancer cell lines viz., HeLa, SiHa and CaSki were transfected with the optimized and non-optimized *CDK1,* and *NUF2* synchronization was performed using double thymidine block. Cell cycle analysis revealed that the non-optimized genes had a slow cell cycle compared to optimized genes, where the cells with optimized genes entered G2 at 10^th^ hr compared to non-optimized. This might suggest that the non-optimized genes help in maintaining proper function of cell cycle dependent genes. Previous studies showed that optimized codons are linked to differentiation while non-optimized are linked to proliferation genes^57–59^.

*CDK1,* and *NUF2* are related to cell cycle and apoptotic regulator mechanism^60–63^. Non-optimized codons help in efficient translation and co-translational protein folding. The mRNA decay rate of genes is also associated with CUB^64,65^. The non-optimized cell-cycle dependent genes have less stability to maintain the efficient functioning and degradation of the genes and proteins. Accumulation of such genes and stability of the genes could be associated with proliferation thereby leading to cancer.

The apoptosis assay was performed to understand the effect of codon optimization on apoptosis using Annexin V-7AAD staining. We observed that cells with optimized genes had more cells in late apoptotic phase than the ones with non-optimized codons. Approximately 70% of *CDK1op,* and *NUFop* had cells in late apoptotic phase than their respective non-optimized codons. This could be attributed to a phenomenon called mitotic catastrophe where cells with shorter cell cycle undergo apoptosis^60,63,66^. Mitotic catastrophe is associated with untimed entry of cells in M phase^67^. This could possibly be one of the mechanisms of the cells to maintain the normal state. The cells with mitotic slippage or shorter cell cycle phase tend to evade the cell cycle checkpoint, which could possibly lead to oncogenesis and therefore, apoptotic pathways for these cells are activated for maintaining the cell cycle regulation^60–62,66–68^.

Essential genes such as housekeeping genes like *GAPDH, ACTB, TUB* use optimized codons^9,38,69^. Genes such as *TP53 and KRAS* use non-optimized codons. Optimization of these genes has led to loss of functions in these genes. Synonymous mutations of *TP53* caused it to lose its tumor suppressor activity, leading to increased cell proliferation^70–72^. However, their roles in apoptosis have not been studied so far. It will be interesting to optimize the codons of essential genes and express it in human cell lines thereby checking its effect on cell death and survival. However, some reports suggest, any mutation in essential genes, can lead to mitotic catastrophe, which is cell death due to biological, physical and chemical stress.

Therefore, cell cycle dependent genes use non-optimal codon usage to maintain gene structure, stability and cellular functions. Perturbation in the codon usage could lead to functional implications following apoptosis of the cells and perturbed cell cycle, which can lead to various cellular and functional disorders.

