## Supplementary Figures for "Non-Optimal Codon Usage Regulates Cell Cycle Progression: Functional Insights Of Codon Optimization of CDK1 and NUF2 Genes"

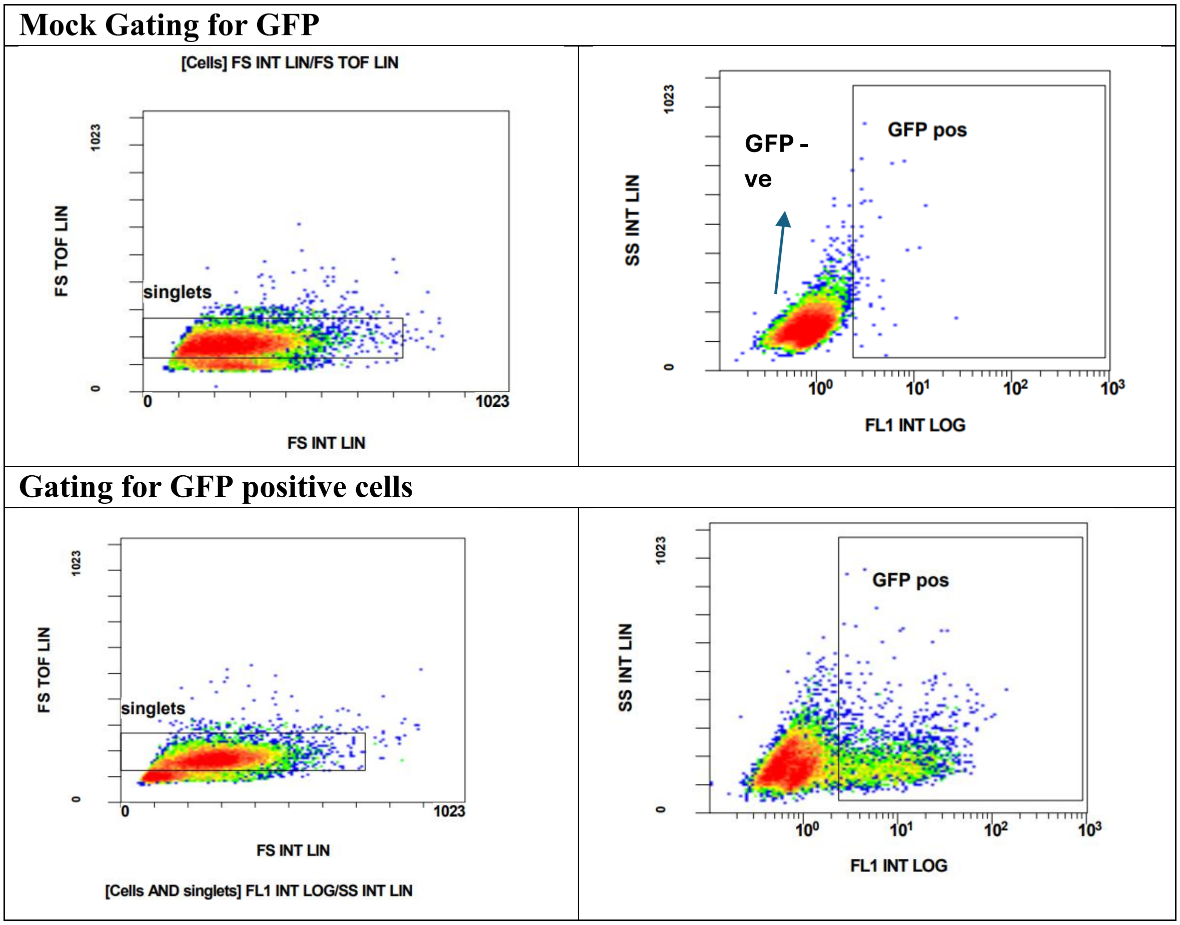


**Figure S1A: Gating of GFP for flow cytometry analysis:**

Left: gating for single cells., Right: gating for GFP cells. Top Right: gating for GFP -ve cells using untransfected cells as mock. Bottom right: using eGFP transfected cells as positive control for GFP+ve.

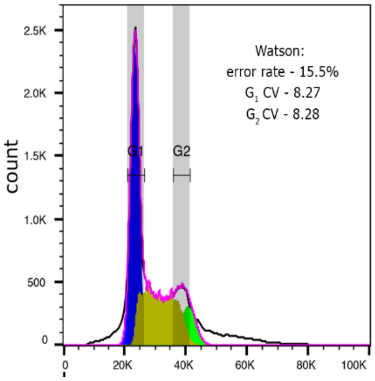


**Figure S1B: cell cycle phase.**

A: Hoechst stains are used to separate G0-G1/S/G2/M. The first peak shows G0-G1, followed by a plateau S and a small peak at G2/M. the linear count of G2/M is twice that of G0-G1.

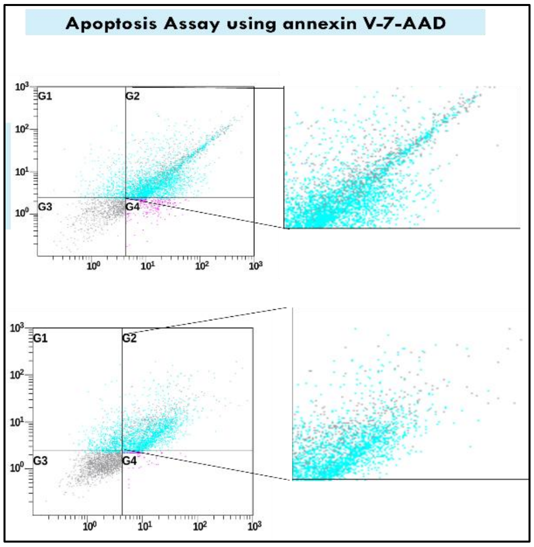


**Figure S2. Flow cytometry analysis for Apoptosis using Annexin V-7AAD:**
(Representative) Flow cytometric analysis of apoptosis assay for CDK and CDKop.,
